# White Matter Connectivity Reflects Success in Musical Improvisation

**DOI:** 10.1101/218024

**Authors:** Tima Zeng, Emily Przysinda, Charles Pfeifer, Cameron Arkin, Psyche Loui

**Author notes:** Corresponding author: Psyche Loui, Department of Psychology, Program in Neuroscience & behavior, Wesleyan University, 207 High Street, Middletown, CT, USA 06459, 1(860)685-2313.

## Abstract

Creativity is the ability to produce work that is novel, high in quality, and appropriate to an audience. One domain of creativity comes from musical improvisation, in which individuals spontaneously create novel auditory-motor sequences that are aesthetically rewarding. Here we test the hypothesis that individual differences in creative behavior are subserved by mesial and lateral differences in white matter connectivity. We compare jazz improvising musicians against classical (non-improvising) musicians and non-musician control subjects in musical performance and diffusion tensor imaging. Subjects improvised on short musical motifs and underwent DTI. Statistical measures of fluency and entropy for musical performances predicted expert ratings of creativity for each performance. Tract-Based Spatial Statistics (TBSS) showed higher Fractional Anisotropy (FA) in the cingulate cortex and corpus callosum in jazz musicians. FA in the cingulate also correlated with entropy. Probabilistic tractography from these mesial regions to lateral seed regions of the arcuate fasciculus, a pathway known to be involved in sound perception and production, showed mesial-to-lateral connectivity that correlated with improvisation training. Results suggest that white matter connectivity between lateral and mesial structures may integrate domain-general and domain-specific components of creativity.

## 1. Introduction

Creativity is the ability to produce work that is original and appropriate to an audience (Hollenberg, 1999), but how the human brain enables creativity is only beginning to be understood. Recent advances in human neuroscience show that the intrinsic structural connectivity of brain networks underlies much of behavior. Variations in white matter microstructure are related to individual differences in specific cognitive skills and are influenced by experience and training (Johansen-Berg, 2010; Lerch et al., 2017; Scholz, Klein, Behrens, & Johansen-Berg, 2009). However, little is known about how these variations in brain anatomy and connectivity relate to creativity, partly because of challenges in behaviorally defining creativity.

Creative tasks differ from standard laboratory tasks in that there is no uniquely correct answer. Studies on creativity have relied on divergent thinking tasks, in which participants generate varied responses to a certain item, and results are scored on fluency, originality, and flexibility (Guilford, 1967). High scores on the divergent thinking task are associated with higher fractional anisotropy in white matter of the frontal lobe, including the corpus callosum and cingulate cortex (Takeuchi et al., 2010). This is consistent with resting state functional MRI studies which found higher functional connectivity in high scorers on the divergent thinking task between the medial prefrontal cortex and posterior cingulate cortex (Takeuchi et al., 2012). These mesial areas constitute core regions of the Default Mode Network, which is usually deactivated during cognitive tasks (Beaty et al., 2014; Greicius, Supekar, Menon, & Dougherty, 2009). Interestingly, those who scored high on divergent thinking tasks showed lower task-induced deactivation in the DMN during a working memory task (Takeuchi et al., 2011), as well as higher resting-state functional connectivity between the lateral inferior prefrontal cortex (specifically right inferior frontal gyrus) and the DMN (Beaty et al., 2014). These results offer the intriguing hypothesis that creativity, as assessed via divergent thinking, could arise from increased connectivity or cross-wiring between mesial and lateral structures.

While divergent thinking tests are a useful laboratory assessment of relatively domain-general creative behavior, creativity researchers have advocated using domain-specific measures to capture the hypothesized integration between domain-general and domain-specific functions that characterizes creative behavior in everyday life (Baer, 1993; Runco, 2008). One specific domain of creativity comes from musical improvisation, such as in jazz music, where musicians spontaneously create novel auditory-motor sequences as a result of long-term practice and domain-specific training (Beaty, 2015; Bengtsson, Csikszentmihalyi, & Ullen, 2007; Limb & Braun, 2008). Previous fMRI work on musical improvisation identified increased activity in prefrontal mesial regions including the medial prefrontal/orbitofrontal cortex and anterior cingulate cortex (Limb & Braun, 2008; Liu et al., 2012), but also activity in the inferior frontal gyrus (IFG) and the superior temporal gyrus (STG), during musical improvisation (Bengtsson et al., 2007; Donnay, Rankin, Lopez-Gonzalez, Jiradejvong, & Limb, 2014). The IFG and STG, classic regions within the language network, are connected by the arcuate fasciculus, a white matter pathway that enables perception-action coupling (Loui, Alsop, & Schlaug, 2009; Pulvermüller & Fadiga, 2010). Long-term training in auditory-motor mapping, such as musical training, is associated with increased volume of the arcuate fasciculus (Halwani, Loui, Rueber, & Schlaug, 2011; Moore, Schaefer, Bastin, Roberts, & Overy, 2014; Oechslin, Imfeld, Loenneker, Meyer, & Jancke, 2010). The arcuate fasciculus is part of the dorsal stream: a structural and functional brain pathway between temporal and frontal lobes (Friederici, Bahlmann, Heim, Schubotz, & Anwander, 2006; Saur et al., 2008) that subserves sensorimotor mapping of sound to articulation, and is crucial for language and music as well as auditory-motor behavior more generally (Hickok & Poeppel, 2007; Loui, 2015).

While the arcuate fasciculus connects the lateral temporal and lateral frontal regions that constitute the auditory perception-action network, it is unclear how these lateral regions might enable the spontaneously novel auditory-motor behavior that characterizes real-time musical creativity as is observed in musical improvisation. Integrating the multiple lines of structural and functional neuroimaging evidence reviewed above regarding divergent thinking, perception-action coupling, and musical improvisation, our hypothesis is that the lateral endpoints of the arcuate fasciculus might show enhanced connectivity with mesial regions among individuals who are better able to create novel auditory-motor sequences. Here we test this hypothesis of enhanced integration between lateral and mesial regions underlying auditory-motor creativity, by comparing Jazz improvising musicians (who routinely generate spontaneous musical sequences as part of their practice) against Classical musicians (who also routinely generate highly trained musical sequences, but without the characteristic spontaneity), against non-musicians (who have little to no training in generating musical auditory-motor sequences). Using diffusion tensor imaging, we first tested the hypothesis that Jazz musicians might have different patterns of Fractional Anisotropy (an index of white matter integrity) compared to Classical musicians and non-musicians. Secondly, we tested whether FA might reflect differences in fluency and entropy, two measures of information content within recorded musical output that we show for the first time to predict expert listeners’ ratings of creativity. Thirdly, we seeded results from whole-brain differences in FA towards regions of interest within the endpoints of the arcuate fasciculus for probabilistic tractography, to assess whether the FA differences that we identify in Jazz musicians are preferentially connected to specific regions in the brain.

## 2. Materials and Methods

### 2.1 Subjects

Fifty-five subjects in total participated in this study. Students were recruited from Wesleyan University and the Hartt School of Music, and received monetary compensation or course credit for their participation. Thirty-nine of these participants had MRI data and fit the criteria for one of the three groups based on their reported musical experience (as reported below), and thus were used in group comparisons. Fifty-one subjects completed the musical improvisation task, but the improvisation data were lost in two subjects due to technical difficulties, leaving forty-nine subjects who had musical improvisation data. Thirty-eight of these subjects were rated by an expert jazz instructor and professional musician. Thirty-nine of the subjects who had musical improvisation data also had MRI data, and thus were used in brain-behavior correlations.

The jazz group (n = 15) was defined by the following criteria: 1) 5+ years of training in music that included improvisation, and 2) Current participation in improvisatory musical activities for 1+ hours per week. The classical group (n = 12) was defined by the following criteria: 1) 5+ years of musical training not including improvisational training. 2) Current participation in musical activities for 1+ hour per week. The non-musician group (n = 12) included participants who had less than 5 years of previous musical training. The three groups were matched in age, general intelligence and working memory, and low-level pitch discrimination ability, and age of onset of musical training (see Procedures). Both the jazz and classical groups had a longer duration of musical training than the non-musicians group, but there was no significant difference between the jazz and classical groups. The jazz group had an average of 5.7 years of training in improvisation specifically, which was significantly higher than both the classical and non-musician groups who were close to 0 years of improvisation training. Subjects from the jazz group reported improvisatory musical activities in instruments including piano (n = 2), guitar (n = 5), saxophone (3), drums (4), clarinet (1). Subjects from the classical group reported non-improvisatory musical activities in guitar (n = 3), piano (n = 1), clarinet (2), flute (1), violin (1), voice (1), horn (1), drums (1), pipa (1). All subjects gave informed consent as approved by the Institutional Review Boards of Wesleyan University and Hartford Hospital.

**Table 1.** Demographics, training information, and baseline behavioral tests in jazz, classical, and non-musician subjects.

|  | **Non-musicians**<br>**(n = 12)** | **Classical Musicians**<br>**(n = 12)** | **Jazz Musicians**<br>**(n = 15)** | **Non-musicians**<br>**vs.**<br>**Classical Musicians** | **Non-musicians**<br>**vs. Jazz Musicians** | **Classical Musicians**<br>**vs Jazz Musicians** |
| --- | --- | --- | --- | --- | --- | --- |
| **Pitch Discrimination (Hz)** | 10 (13) | 6.0 (3.1) | 4.7 (3.1) | ns | ns | ns |
| **Digit Span (digits)** | 7.1 (1.6) | 7.3 (1.8) | 8.0 (1.5) | ns | ns | ns |
| **Shipley (raw score)** | 17.1 (1.8) | 17.3 (1.7) | 17.1 (1.8) | ns | ns | ns |
| **Age (years)** | 19.8 (.97) | 21.8 (4.8) | 20.7 (1.4) | ns | ns | ns |
| **Age of onset (years)** | 9.2 (2.4) | 7.6 (2.4) | 8.3 (2.7) | ns | ns | ns |
| **Duration of training (years)** | 2 (1.5) | 10.3 (1.9) | 8.7 (3.3) | t(22) = 11.1<br>p < .001 | t(25) = 6.14<br>p < .001 | ns |
| **Duration of improv training (years)** | 0 (0) | 1 (1.7) | 5.7 (3.1) | ns | t(25) = 5.9<br>p < .001 | t(25) = 4.7<br>p < .001 |

### 2.2 Procedures.

After subjects gave informed consent, they completed the baseline tasks for the experiment including a pitch discrimination threshold-finding test to rule out differences in pitch perception (Loui, Guenther, Mathys, & Schlaug, 2008), the Shipley Institute of Living Scale to rule out possible intellectual impairment (Shipley, 1940), and a digit span task that tested working memory (Baddeley, 2003). None of these tasks showed significant differences between the three groups (Table 1). Subjects then completed a questionnaire on their musical background including questions about the age of onset and duration of general musical training, the duration of jazz and improvisation training, the amount of time spent on musical activities, and a self-rating of their improvisation skills. The baseline tasks were followed by an improvisation continuation task that asked participants to improvise based on simple musical phrases. All participants completed this task on the keyboard; jazz and classical musicians who reported other instruments additionally completed the task on their own instrument. In a separate session participants were transported to the Hartford Institute of Living where the MRI took place.

#### 2.2.1 Improvisation Continuation Task.

Subjects were given 12 musical motifs (short musical fragments) as inspiration to improvise. Each motif consisted of one measure, which lasted 2.4 seconds (4 beats at 100 beats per minute) centering around MIDI pitch C4, that varied in rhythm and/or pitch. (See https://wesfiles.wesleyan.edu/home/ploui/web/JazzCreativity/ImprovCont/Motives/ for audio files of all 12 motifs.) For each trial subjects listened to one motif twice, then continued the motif by repeating it and then improvising on it for a total of 16 measures. There was a constant metronome in the background throughout the trial to keep time. Subjects were instructed to play as creatively and imaginatively as possible. This task was designed to be sensitive to each player’s musical improvisation ability, while being as accessible as possible across musicians and non-musicians. Responses were collected on a Casio PX150 MIDI keyboard that was connected to Max/MSP (Zicarelli, 1998), a software environment which in which custom-made software recorded the keystrokes and exported the MIDI data as text files for further analysis. All performances were also video and audio recorded using a Zoom Q8 video camera.

#### 2.2.2 Magnetic Resonance Imaging Acquisition.

High-resolution T1 and DTI images were acquired in a 3T Siemens Skyra MRI scanner at the Olin Neuropsychiatry Research Center at the Institute of Living. The anatomical images were acquired using a T1-weighted, 3D, magnetization-prepared, rapid-acquisition, gradient echo (MPRAGE) volume acquisition with a voxel resolution of 0.8 × 0.8 × 0.8 mm^3^. The diffusion images were acquired using a diffusion-weighted, spin-echo, echo-planar imaging sequence (TR = 4.77 s, voxel size = 2.0 × 2.0 × 2.0 mm^3^, axial acquisition, 64 noncollinear directions with b-value of 1000 s/mm^2^, 64 noncollinear directions with b-value of 2000 s/mm^2^, 1 image with b-value of 0 s/mm^2^).

### 2.3 Data Analysis

#### 2.3.1 Improvisation Continuation Task

Each subject’s performed output on the keyboard was exported as MIDI data from Max/MSP to Matlab for objective analysis. Information-theoretic analyses yielded two measures: fluency and entropy, that reflect the information content of musical recordings (Hansen & Pearce, 2014; Lartillot & Toiviainen, 2007). Fluency is the total number of notes produced for each trial. It is similar to fluency as the number of responses in domain-general creativity tasks such as the classic Torrance Test of Creative Thinking (Torrance, 1968). Entropy is a measure of information content (Shannon, 1948). It is defined as H(X) = −Σp_i_*log(p_i_), where p_i_ = probability of each note of each subject’s output in each trial. Shannon entropy is analogous to the score for originality in the Torrance test, where more unusual (low probability) responses are given more points. Fluency and entropy values were averaged across the 12 trials for each subject.

Additionally, two independent professional jazz musicians and music instructors were recruited as expert blind raters to listen to the audio recording of each trial and to rate them for creativity. One rater completed 120 ratings (10 subjects * 12 trials each). The second rater completed 380 ratings (38 subjects * 12 trials each). The two raters’ ratings for the first 120 recordings were highly correlated (r(118) = .544, p < .001). Ratings from the second rater were used for further analysis.

#### 2.3.2 DTI analysis.

Voxelwise statistical analysis of the FA data was carried out using TBSS (Tract-Based Spatial Statistics (Smith et al., 2006)), part of FSL (Smith et al., 2004). TBSS projects all subjects' FA data onto a mean FA tract skeleton, before applying voxelwise cross-subject statistics. First, FA images were created by fitting a tensor model to the raw diffusion data using FDT, and then brain-extracted using BET (Smith, 2002). All subjects' FA data were then aligned into a common space using the nonlinear registration tool FNIRT (Andersson, Jenkinson, & Smith, 2007a, 2007b), which uses a b-spline representation of the registration warp field (Rueckert et al., 1999). Next, the mean FA image was generated and thinned to create a mean FA skeleton which represents the centers of all tracts common to the group. Each subject's aligned FA data was then projected onto this skeleton and the resulting data fed into voxelwise cross-subject statistics. In a first contrast, jazz musicians were compared against classical musicians and non-musicians. Classical musicians were then compared against jazz and non-musicians, and non-musicians were compared against the two musician groups.

Next, the FA skeleton was correlated with fluency and entropy, variables obtained from each individual subject’s behavioral data. Each subject’s mean entropy and fluency was entered into a design matrix and a contrast matrix. Permutation tests were carried out using Randomise with the TFCE method in FSL to identify positive correlations between FA and each behavioral variable (Smith & Nichols, 2009).

In addition to TBSS, probabilistic tractography was applied to identify the dorsal and ventral arcuate fasciculus in each hemisphere of each brain from regions of interest in the temporal lobe and frontal lobe. ROIs were hand drawn over the FSL 1mm FA template on the left and right inferior frontal gyrus (IFG), superior temporal gyrus (STG), and middle temporal gyrus (MTG). Each ROI was then warped to each participant’s native space FA image. Average and standard deviations of volumes and xyz coordinates for the ROIs can be seen in table 2. There were no main effects of group for the volume, locations, or FA of the projected ROIs, and no difference in whole-brain volume or FA between groups (all p > .05). Probabilistic tractography was done using IFG as seed region and STG or MTG as waypoint mask, then the reverse STG or MTG as seed region and IFG as waypoint mask. Each resultant tract was averaged and then thresholded at 10% of its robust intensity level to minimize extraneous tracts. Consistent with previous studies (Halwani et al., 2011; Loui et al., 2009; Loui, Li, & Schlaug, 2011), tracts identified between STG and IFG were labeled as dorsal arcuate fasciculus, whereas tracts identified between MTG and IFG were labeled as ventral arcuate fasciculus. Tract volume and FA were exported for statistical comparisons. Additionally, to enable visualization all subjects' tracts and FA images were aligned and normalized to the FSL 1 mm FA template using both linear registration (FLIRT) (Jenkinson & Smith, 2001) and nonlinear registration tools (FNIRT), and canonical tract images were created by averaging each binarized tract across subjects in each group, and thresholding voxels below the median. Thus, for each group a voxel was considered part of the canonical tract if more than 50% of the group had an identified tract in that voxel.

**Table 2.**
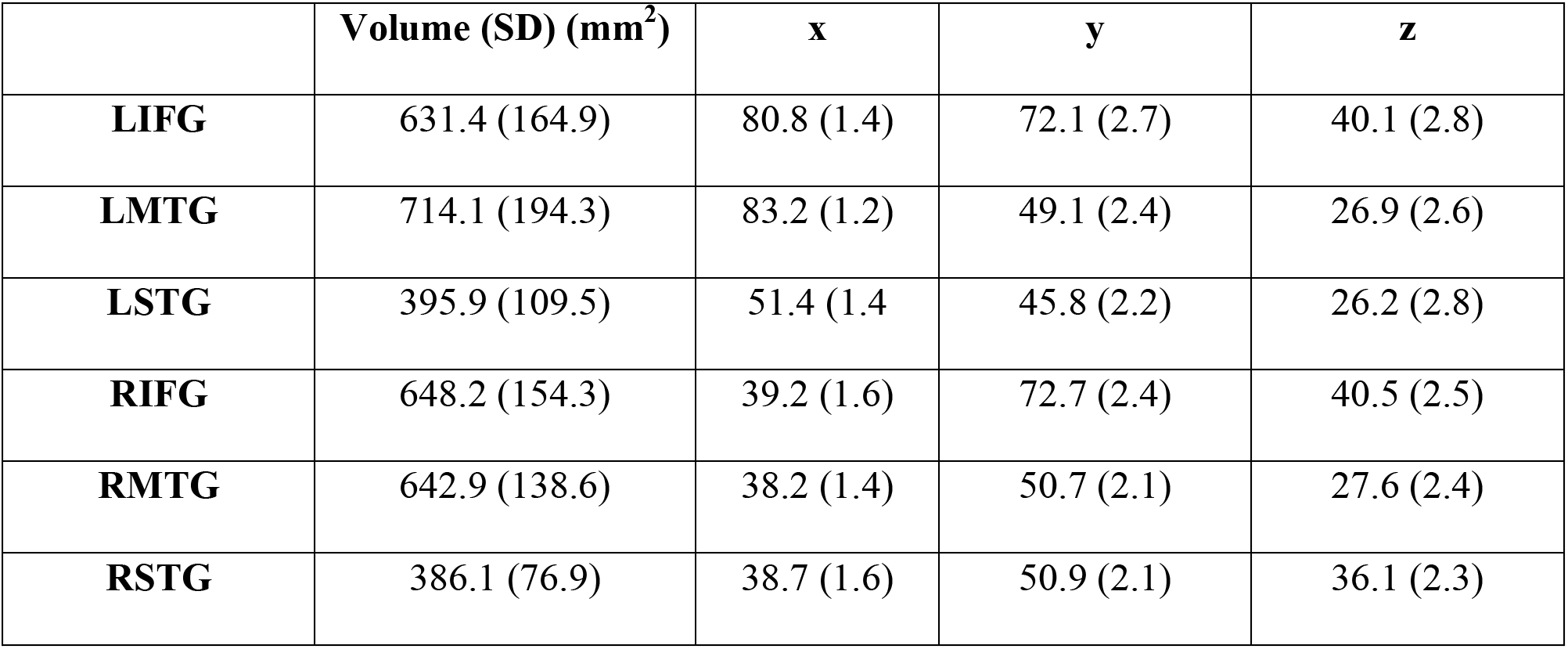
Mean and SD of ROI volumes and locations.

|  | **Volume (SD) (mm<sup>2</sup>)** | **x** | **y** | **z** |
| --- | --- | --- | --- | --- |
| **LIFG** | 631.4 (164.9) | 80.8 (1.4) | 72.1 (2.7) | 40.1 (2.8) |
| **LMTG** | 714.1 (194.3) | 83.2 (1.2) | 49.1 (2.4) | 26.9 (2.6) |
| **LSTG** | 395.9 (109.5) | 51.4 (1.4) | 45.8 (2.2) | 26.2 (2.8) |
| **RIFG** | 648.2 (154.3) | 39.2 (1.6) | 72.7 (2.4) | 40.5 (2.5) |
| **RMTG** | 642.9 (138.6) | 38.2 (1.4) | 50.7 (2.1) | 27.6 (2.4) |
| **RSTG** | 386.1 (76.9) | 38.7 (1.6) | 50.9 (2.1) | 36.1 (2.3) |

To test the specific hypothesis of increased structural connectivity between mesial and lateral regions in jazz improvising musicians, the TFCE-corrected t-statistic image from a positive TBSS group comparison (contrasting jazz musicians against classical musicians and non-musicians) was then thresholded at .95 to include only the 1196 voxels that were significant at the p < .05 TFCE-corrected level. This significant cluster (labeled CC to denote its location in the Cingulate Cortex and Corpus Callosum) was then warped to the native FA image of each subject, and used as a seed ROI towards the separate waypoint masks of STG, MTG, and IFG of each hemisphere. The STG, MTG, and IFG were also used as separate seed ROIs towards the CC as waypoint mask, and each resultant tract was again individually averaged and thresholded, exported, and averaged across subjects using similar procedures as for the identification of the arcuate fasciculus mentioned above.

## 3. Results

### 3.1 Behavioral results: Objective measures predict expert ratings of creativity.

For the Improvisation Continuation task, expert ratings of creativity were significantly higher for the jazz group compared to the classical and non-musician groups (F(2,35) = 12.8, p < .001; see Figure 1). Objective measures of performance showed higher fluency and higher entropy for the jazz group compared to the other two groups (fluency: F(2,46) = 11.5, p < .001 ; entropy: F(2,46) = 7.3, p = .001; see Fig. 1), confirming more fluent production (higher number of notes produced) and higher information content from behavioral output of jazz players. Subjective ratings of creativity by the expert were highly correlated with objective measures of fluency (r(33) = .74, p < .01) and entropy (r(33) = .74, p < .01). In addition, fluency was correlated with duration of general musical training (r(48) = .33, p = .022), duration of improvisation training (r(48) = .36, p = .01), and hours per week of reported improvisatory practice (r(48) = .38, p = .008). Entropy was correlated with duration of improvisational training (r(48) = .30, p = .036), and hours per week of reported improvisatory practice (r(48) = .29, p = .046). Expert ratings of creativity were correlated with duration of improvisational training (r(37) = .50, p = .001), and reported hours per week of improvisatory practice (r(37) = .50, p = .001) but not with duration of general musical training. These results confirm that musicians with jazz improvisation training are perceived by experts as more creative, and that objective measures derived from musical output can accurately predict expert subjective ratings of creativity.

**Figure 1.**
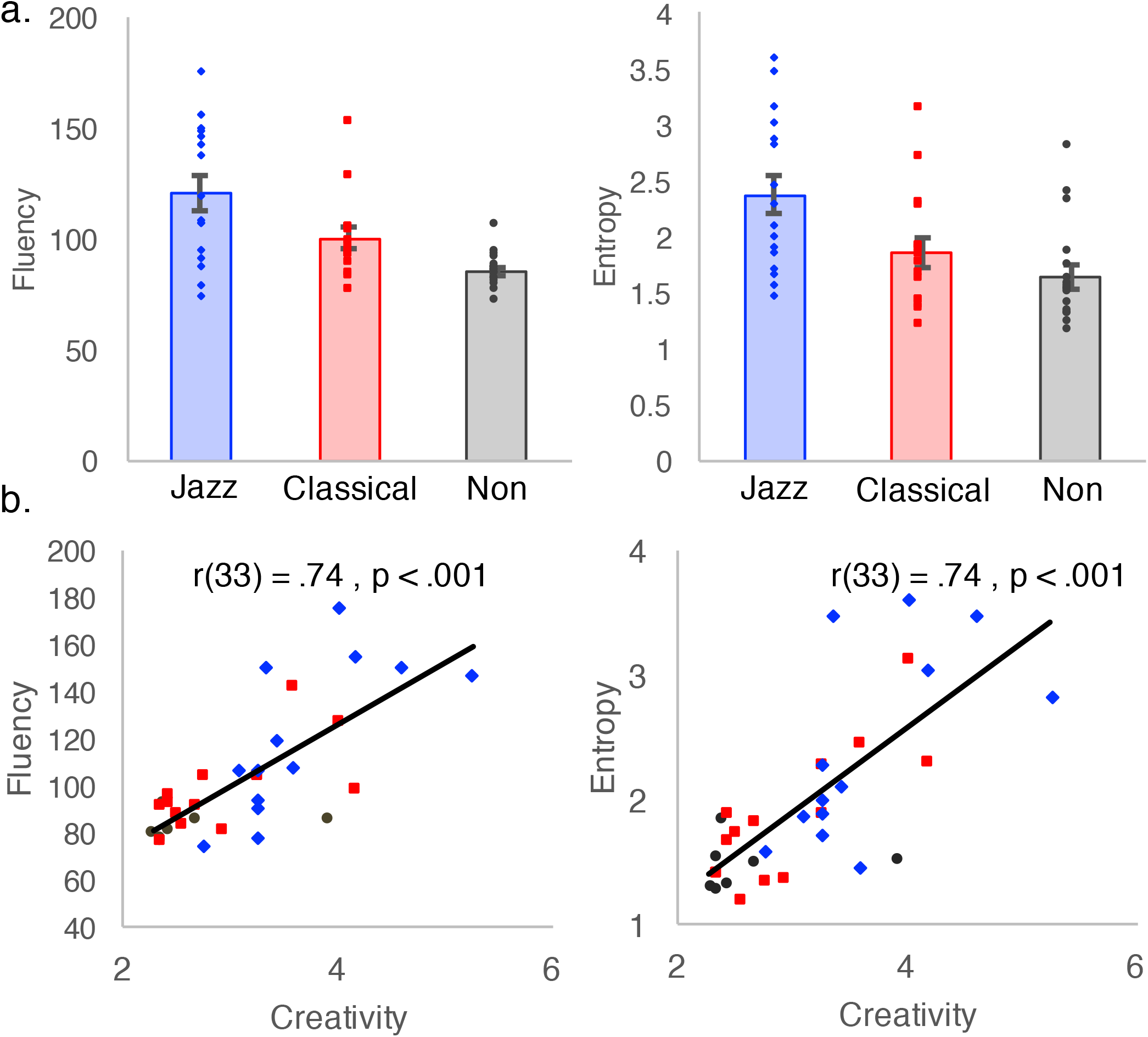
**a.** Fluency and entropy scores for jazz musicians, classical musicians, and nonmusicians. **b.** Measures of fluency and entropy predict expert ratings of creativity. Diamonds = jazz musicians; squares = classical musicians; circles = non-musicians.

### 3.2 TBSS: higher FA in mesial structures.

A TBSS whole-brain comparison showed that the jazz group had a higher FA in a region spanning the corpus callosum and anterior cingulate (1196 voxels, centered on x = −11, y = 5, z = 23, p < .05 TFCE-corrected). This same cluster, hereafter labeled CC (cingulate cortex/corpus callosum), shows higher FA in jazz musicians compared to the classical musician group (t(25) = 3.2, p = .004) and the non-musician group (t(25) = 4.4, p < .001) (see Figure 2a). FA in the CC is also significantly correlated with duration of improvisation training (r(38) = .48, p = .002) and number of hours of reported improvisatory practice per week (r(38) = .61, p <.001).

**Figure 2.**
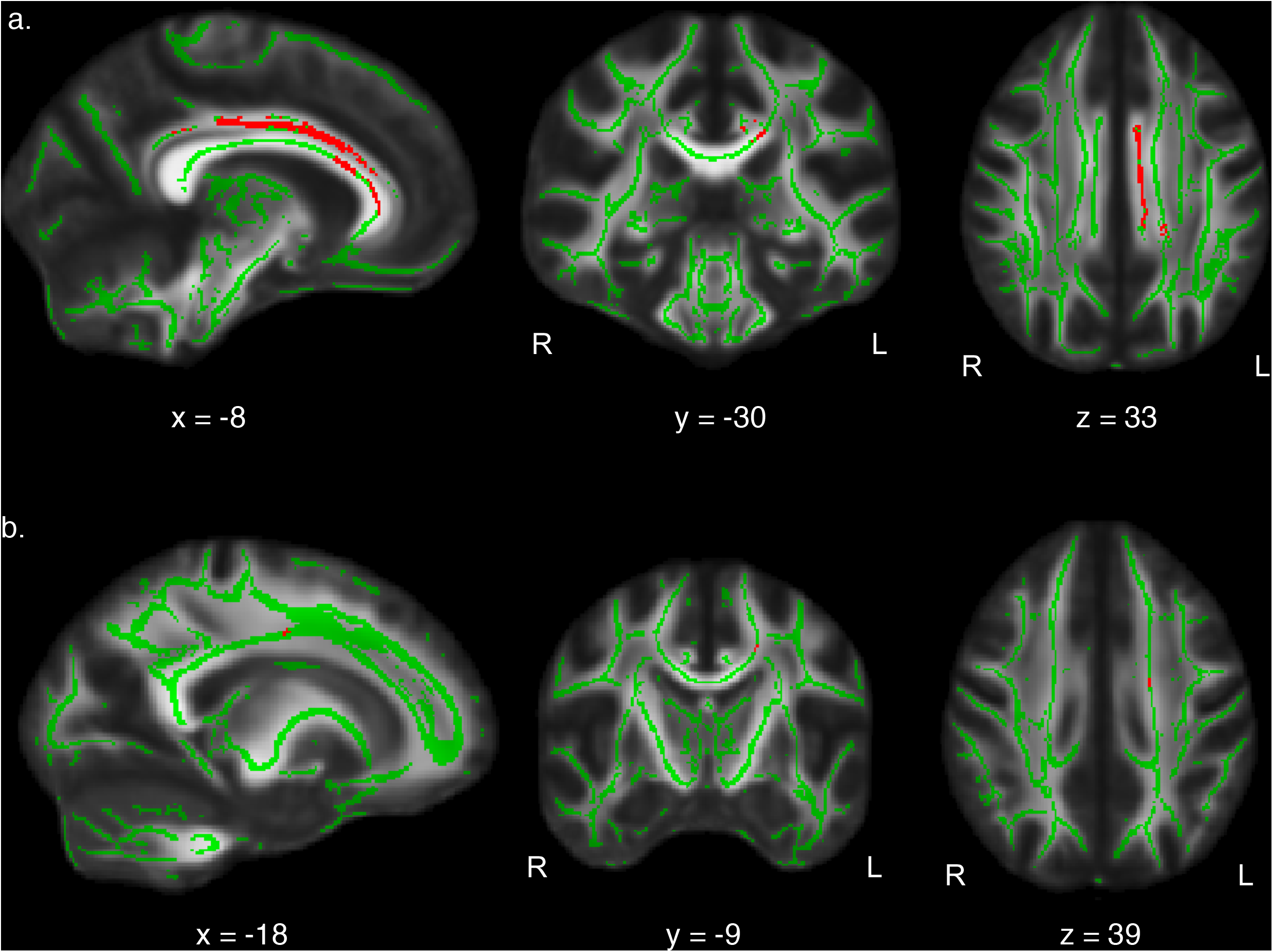
**a.** TBSS result showing higher FA in the jazz group compared to all classical musicians and nonmusicians. Mean FA skeleton across all subjects is shown in green over the FMRIB58_FA_1mm template. A cluster of 1196 voxels with higher FA in the jazz group (p < .05 TFCE-corrected) is shown in red. **b.** TBSS correlation results showing a cluster of 37 voxels (red) that are correlated with entropy (p < .05 TFCE-corrected) on the mean FA skeleton across all subjects (green) over the FMRIB58_FA_1mm template.

### 3.3 Whole-brain correlations with entropy.

A whole-brain TBSS correlation between FA and the behavioral measure of entropy showed a significant cluster (37 voxels centered on x = −18, y = −9, z = 39, p < .05 TFCE-corrected) in the left cingulate cortex (see Figure 2b). Whole-brain TBSS correlation between FA and fluency yielded no significant voxels at the p < .05 TFCE-corrected level.

### 3.4 Arcuate fasciculus

As the arcuate fasciculus is part of a perception-action network that has previously been associated with musical aptitude (Loui et al., 2009) and musical training (Halwani et al., 2011), ROI-based probabilistic tractography was used to identify the four branches of the arcuate fasciculus (left dorsal, left ventral, right dorsal, and right ventral). To correct for multiple comparisons for each of the four branches, a Bonferroni-corrected p-value of p = .0125 was used as the threshold for significant differences. No significant main effect of group or correlations with behavioral measures were observed in either branch using this criterion; however there was a significant correlation between volume of the left dorsal arcuate fasciculus, identified as tracts between left STG and left IFG, and duration of improvisation training (r(38) = .429, p = .006) and number of hours of reported improvisatory practice per week (r(38) = .391, p = .014), thus providing partial support for differences in the perception-action network that are related to improvisation training.

### 3.5 Mesial and lateral structures.

Results from TBSS had identified robust FA differences in mesial structures related to musical improvisation, and partial support for differences in the lateral perception-action network. To investigate whether mesial structures are preferentially connected to specific lateral regions in the brain in jazz musicians, the mesial CC (the significant cluster obtained from the TBSS group comparison, which included voxels in the corpus callosum and anterior cingulate), was then used as a region of interest for probabilistic tractography to investigate its patterns of connectivity to the endpoints of the arcuate fasciculus (left and right IFG, STG, and MTG). No significant differences were observed for tracts between the MTG and significant cluster on either hemisphere, but significant differences were observed in tracts between the CC and the right IFG and the left STG, described below.

#### 3.5.1 IFG to CC Tracts.

For tracts identified between CC and RIFG, a main effect of group on FA was significant (F(2,36) = 5.0, p = .012), where jazz musicians had the highest FA, followed by classical musicians and then non-musicians (jazz vs. classical: t(25) = 2.0, p = .054, jazz vs. non: t(25) = 2.8, p < .01). FA of the same tract was also correlated with measures of entropy (r(32) = .4, p = .022) and fluency (r(32) = .37, p = .034), and also with the duration of jazz training (r(38) = .37, p = .022) and reported number of improvisation practice hours per week (r(38) = .411, p = .009), but not with duration or age of onset of general musical training. Follow-up analyses show a main effect of group on mean diffusivity (F(2,36) = 4.7, p = .015) and radial diffusivity (F(2,36) = 5.6, p = .008), where jazz musicians have the lowest MD and RD (jazz vs. classical, MD: t(26) = 1.7, p = .097, RD: t(26) = 1.9, p = .06; jazz vs. non-musician, MD: t(26) = 2.9, p = .007, RD: t(26) = 3.0, p = .005). MD and RD are negatively correlated with duration of improvisation training (MD: r(39) = -.412, p = .009, RD: r(39) = -.383, p = .016). There were no significant differences in tract FA between CC and the LIFG.

#### 3.5.2 STG to CC Tracts

For tracts identified between LSTG and CC, a main effect of group on FA was significant (F(2,36) = 4.6, p = .016). Jazz musicians had the highest FA, followed by classical musicians and then non-musicians (jazz vs. classical: t(25) = 2.1, p = .043; jazz vs. non-musicians: t(25) = 2.7, p = .012). FA of the same tract was also correlated with entropy (r(32) = .36, p = .041) and with the duration of jazz training (r(38) = .37, p = .021) and reported number of hours of improvisatory practice per week (r(38) = .37, p = .021), but not with duration or age of onset of general musical training. Follow-up analyses for the LSTG to CC tract show a main effect of group on mean diffusivity (F(2,36) = 5.1, p = .011) and radial diffusivity (F(2,36) = 5.2, p = .01), where jazz musicians have lower mean and radial diffusivity compared to non-musicians (MD: t(25) = 3.1, p = .005, RD: t(25) = 3.1, p = .005). There was a marginally significant main effect of axial diffusivity (F(2,36) 3.1, p = .056) and follow up t-tests show that the jazz and classical musician groups each have higher AD compared to the non-musician group (jazz vs. non-musicians: t(26) = 2.1, p = .043; classical vs. non-musicians: t(26) =2.2, p = .035). MD and RD are negatively correlated with duration of improvisation training (MD: r(39) = -.333, p = .038, RD: r(39) = -.318, p = .049). There were no significant differences in tract FA between the CC and the RSTG.

**Figure 3.**
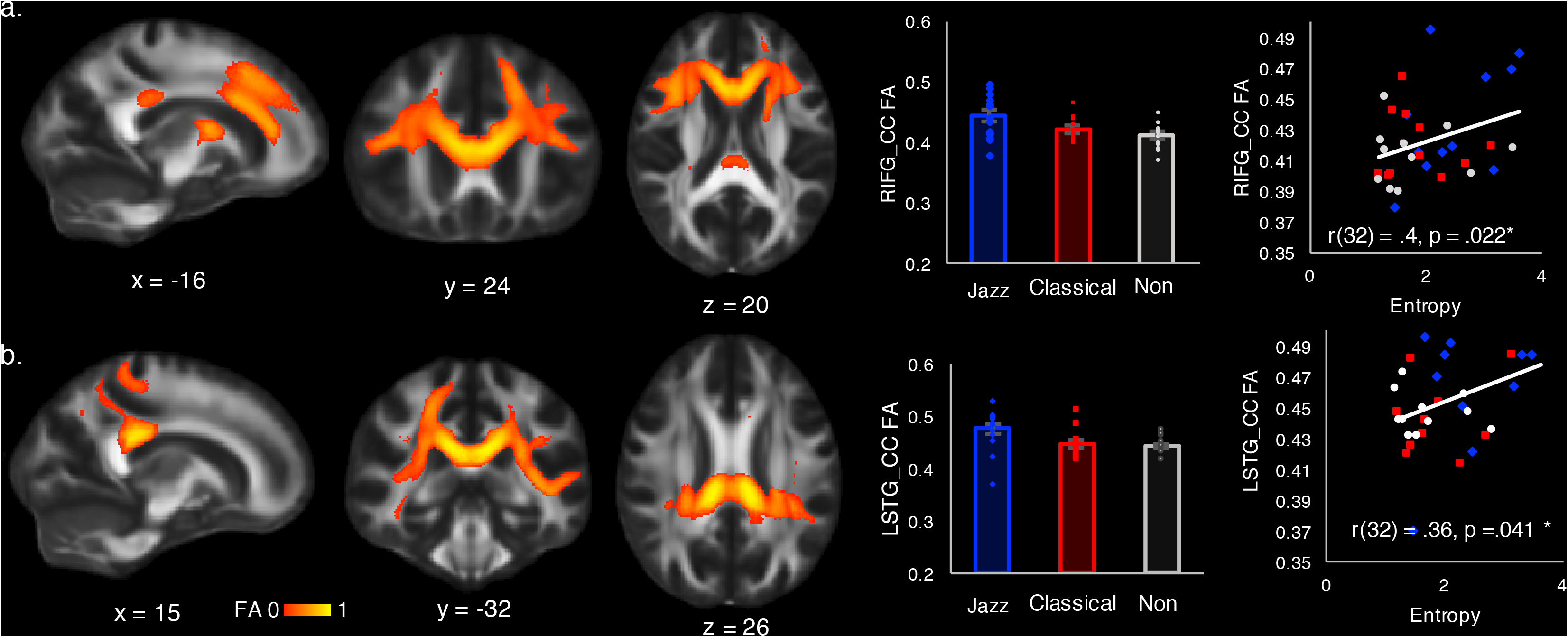
**a.** Tracts identified by probabilistic tractography between right IFG and TBSS-identified significant cluster (shown in red in Fig. 2a). This tract shows highest mean FA in jazz musicians. Mean FA of this tract is also correlated with duration and intensity of training as well as entropy of behavioral output (blue = jazz musicians; red = classical musicians; black = non-musicians). **b.** Tracts identified by probabilistic tractography between left STG and the same TBSS-identified significant cluster. This tract shows highest mean FA in jazz musicians. Mean FA of this tract is also correlated with duration and intensity of training as well as entropy of behavioral output (blue = jazz musicians; red = classical musicians; black = non-musicians).

## 4. Discussion

White matter connectivity predicts individual differences in creative behavior, specifically in musical improvisation. By combining diffusion tensor imaging and subjective and objective analyses of creative output, we show that individual differences in brain connectivity are predictive of quantifiable creative behavior within one’s domain of expertise. Results relate brain connectivity to creative output, and have implications for the understanding of brain structure especially as it relates to training and to the fostering of positive traits such as creativity.

Behavioral data confirm that jazz musicians perform better in our improvisation continuation task both in subjective measures of creativity by external raters and objective measures of fluency and entropy. The high correlation between these subjective and objective measures further supports the use of information theory as an index for creativity in musical improvisation, and offers a novel operational definition of creativity as the fluent production of high information content. While all of these measures were correlated with duration (number of years) and intensity (practice hours) of improvisation training, fluency is also correlated with general music training whereas entropy and creativity are not. This suggests that fluency may reflect general, non-improvisatory musical training that facilitates proficiency on an instrument, whereas entropy and creativity are enhanced more through training in improvisation. Results are not explained by overall duration of musical training, as jazz and classical musicians are similar in this regard.

Low-level differences in perceptual ability and general intellectual functioning and working memory differences also cannot account for these results, as they are similar among the three groups. Differences in technical proficiency at the piano are also unlikely to explain the observed differences in fluency, entropy, and ratings of creativity, as the subjects report training on a variety of instruments, and only a minority of subjects (2 from the jazz group and 1 from the classical group) are pianists. Rather, results are strongly associated with the duration and intensity of improvisation training, quantified here as number of years and hours of practice of improvisation.

Jazz musicians show higher FA, a measure of white matter integrity, in the cingulate cortex and corpus callosum. FA in this region is correlated with improvisational training but not with general music training. Results are also not explained by general differences in cognitive ability or working memory capacity or differences in perceptual acuity, as these are controlled between groups. FA of the cingulate cortex was additionally correlated with measures of entropy. These findings of increased white matter integrity in the corpus callosum and cingulate cortex are consistent with prior work showing associations between the same mesial structures with domain-general measures of creativity, specifically with scores on the divergent thinking test (Takeuchi et al., 2010). Thus, brain structures that underlie divergent thinking, which entails generating original and appropriate ideas, may also underlie musical improvisation, which is the task of generating original and appropriate music ideas in real time. The two tasks, while differing in the form of response, may share some common cognitive demands. Such common cognitive demands may constitute the domain-general components of creativity. These may include idea generation and evaluation, which requires cognitive control and expectancy processing as discussed below.

Increased white matter connectivity in the cingulate cortex could enable greater integration between the cingulate and other regions of the brain. Since musical improvisation, like many acts of spontaneous creativity, relies on the blind variation and selective retention of domain-specific ideas (Campbell, 1960; Jung, Mead, Carrasco, & Flores, 2013), this rapid evaluation of multiple ideas and their associated rewards is in accordance with previous studies that identify the cingulate cortex’s role in reward expectancy. Animal work has shown that neurons in the cingulate cortex are sensitive to reward expectancy for multiple possible outcomes (Hayden, Pearson, & Platt, 2009). fMRI work with humans have found that the dorsal anterior cingulate cortex processes expectancy violations while the ventral anterior cingulate is more influenced by social feedback (Shidara & Richmond, 2002; Somerville, Heatherton, & Kelley, 2006). The dorsal anterior cingulate is also thought to determine the expected value of cognitive control, and thus to assess the costs and benefits of allocating certain cognitive resources (Shenhav, Cohen & Botvinick 2016). These studies together implicate the role of the cingulate cortex in processing expectancy, which dovetails with our recent results showing that jazz musicians process violations in musical expectancy with enhanced early sensitivity and reduced late feedback-dependency (Loui et al., 2016; Przysinda, Zeng, Maves, Arkin, & Loui), which are consistent with these roles of the cingulate cortex.

White matter connectivity in the corpus callosum enables increased integration or communication between the left and right hemispheres, such as between a lateralized region in one hemisphere and its contralateral homolog. Although larger corpus callosum size has been observed in musicians (Schlaug, Jancke, Huang, Staiger, & Steinmetz, 1995), this is the first report of corpus callosum being related to a specific type of musical training. Classic work from corpus callosotomy patients has shown that the corpus callosum is necessary for integrating perceptual information and productive mechanisms between the two hemispheres to enable successful linguistic communication (Gazzaniga & Sperry, 1967). Existing theories on hemispheric asymmetry posit that the left and right hemispheres subserve local vs. global processing in attention (Ivry & Robertson, 1997). Our finding of increased FA in jazz musicians is compatible with both functional accounts of the corpus callosum, as musical improvisation requires integrating perception, ideation, and sound generation, as well as the fluid switching between attention to local and global levels of musical processing (Justus & List, 2005).

Although there is no significant between-group difference in the arcuate fasciculus, the dorsal branch of the left arcuate fasciculus is correlated with duration and intensity of training. This result is consistent with prior reports of the arcuate fasciculus as part of the perception-action pathway that is sensitive to the acquisition of auditor-motor contingencies such as music and language (Halwani et al., 2011; Moore et al., 2014; Qi, Han, Garel, San Chen, & Gabrieli, 2015; Saygin et al., 2013). Importantly, jazz musicians have higher FA in white matter connections between the mesial structures, the cingulate cortex and corpus callosum, and selective endpoints of the arcuate fasciculus. Specifically, there is increased integrity in white matter connections between CC and left STG and right IFG in jazz musicians, and these connections are associated with behavioral measures of musical creativity. Strikingly, these effects are only found in connections to the left STG and right IFG, and not to the right STG, left IFG, or the left or right MTG. Thus, the mesial structures are not generally more connected to the whole brain, but show selective preferential connectivity patterns towards right frontal and left superior temporal regions. The right IFG plays a key role in auditory perception and action; its structural connectivity is linked to abilities in pitch perception and production as well as in tone language acquisition (Loui et al., 2009; Qi et al., 2015). Functional roles of the right IFG include the integration of musical action and perception (Bianco et al., 2016), as well as the successful formation of expectancy in cognitive control tasks (Behan, Stone, & Garavan, 2015). Our results are consistent with the role of RIFG in the musical perception-action network, but also extend existing findings by showing a functional significance of structural connectivity from the RIFG to mesial structures including the cingulate and corpus callosum.

The left STG is part of the classic language network that includes dorsal and ventral pathways from the classic Wernicke’s to Broca’s areas (Friederici, 2009; Hickok & Poeppel, 2007; Parker et al., 2005). Structural connectivity from the left STG via the arcuate fasciculus, part of the dorsal pathway for language, is more left-lateralized in trained singers (Halwani et al., 2011).

The left STG is larger in musicians with absolute pitch (Schlaug, Jancke, Huang, & Steinmetz, 1995). Structural connectivity within the left temporal lobe, specifically between left STG and left MTG, is also enhanced in absolute pitch musicians (Loui, Li, Hohmann, & Schlaug, 2011), and the LSTG is highly functionally connected to multiple areas throughout the brain during music listening especially among individuals with absolute pitch (Loui, Zamm, & Schlaug, 2012). Together these results suggest that LSTG is a hub within an experience-dependent shared network for language and music, that enables the identification of auditory features including the absolute identification of pitch. Although none of the subjects in our study reported having absolute pitch, jazz improvisation training could develop a heightened tonal memory for pitch (Ross, Gore, & Marks, 2005) that engages the same neural mechanisms that absolute pitch musicians possess. By identifying connections between LSTG and mesial structures, the present findings extend previous results by proposing a mechanism through which a relatively domain-specific region for pitch identification may be linked to creativity and expectancy in a relatively domain-general manner. Although mesial structures and lateral structures (such as the language network) are classically thought of as separate systems, our results show that these networks are interconnected.

Additional findings show that jazz musicians have lower mean and radial diffusivity, but no difference in axial diffusivity, in the mesial-lateral connections. As increased RD has been linked to demyelination (Song et al., 2002), lower RD may suggest higher myelination in jazz musicians, but could also be influenced by other factors such as the crossing of multiple fibers (Wheeler-Kingshott & Cercignani, 2009). Lower MD has been observed as a result of spatial learning and memory training (Sagi et al., 2012), and provides evidence for rapid training-induced neuroplastic changes, possibly due to changes to water content in astrocytes (Johansen-Berg, Baptista, & Thomas, 2012). The MD and RD differences observed here are significantly correlated with duration of training and intensity of improvisation practice, but not with general musical training. Together these microstructural findings may suggest that musical improvisation training induces changes in glial cell activity that regulates synaptic plasticity, such as astrocyte water content and myelination. Although this is an intriguing hypothesis, since the current study is a cross-sectional study, we cannot draw any conclusion about the causal relationship between improvisation training and brain connectivity. The differences in brain connectivity may be a result of improvisation training, or they may underlie the subjects’ potential to become experts in improvisation. Further studies that employ a longitudinal design are needed to strengthen this hypothesis. Nevertheless, the current findings, by showing the relation between white matter integrity and jazz improvisation training, add to previous evidence that variations of white matter architecture are related to individual differences in specific cognitive skills (Johansen-Berg, 2010), but extend the findings towards white matter substrates of creative and positive behavior.

## 5. Conclusions

Taken together, results suggest that multiple systems integrate to subserve the creative process. Brain structures that enable improvisational creativity likely include domain-specific as well as domain-general components. Domain-specific components include perception and action as subserved by specific endpoints of the arcuate fasciculus, which is a relatively domain-specific network that pertains to music and language. Domain-general components include the corpus callosum and anterior cingulate, which are sensitive to expectancy and cognitive control. By combining behavioral informatics with diffusion tensor imaging, we detect individual differences in brain connectivity that reflect individual differences in creative behavior. Results may have clinical applications by informing the design of creative music therapy programs as a form of neurorehabilitation for those recovering from neurological and/or communication disorders (Altenmüller & Schlaug; Bringas et al., 2015; Francois, Grau-Sanchez, Duarte, & Rodriguez-Fornells, 2015; Harrison, McNeely, & Earhart, 2017; Ripolles et al., 2015). Most generally, these results may be informative about what creativity entails, how the human brain enables creativity, and how we can learn to be more creative.

## 6. Acknowledgements

We acknowledge Pheeroan AkLaff and John Baboian for serving as expert raters in this study. This research was supported by grant RFP-15-15 from the Imagination Institute (http://www.imagination-institute.org), funded by the John Templeton Foundation. The opinions expressed in this publication are those of the authors and do not necessarily reflect the views of the Imagination Institute or the John Templeton Foundation.

